# The sodium leak channel complex is modulated by voltage and extracellular calcium

**DOI:** 10.1101/740456

**Authors:** Han Chow Chua, Matthias Wulf, Claudia Weidling, Lise Pilgaard Rasmussen, Stephan Alexander Pless

## Abstract

The sodium leak channel (NALCN) is essential for survival in mammals: NALCN mutations are life-threatening in humans and knockout is lethal in mice. However, the basic functional and pharmacological properties of NALCN have remained elusive. Here, we found that the robust function of NALCN in heterologous systems requires co-expression of UNC79, UNC80 and FAM155A. The resulting NALCN channel complex is constitutively active, conducts monovalent cations but is blocked by physiological concentrations of extracellular divalent cations. Our data support the notion that NALCN is directly responsible for the increased excitability observed in a variety of neurons in reduced extracellular Ca^2+^. Despite the smaller number of voltage-sensing residues in the putative voltage sensors of NALCN, the channel complex shows voltage-dependent modulation of the constitutive current, suggesting that voltage-sensing domains can give rise to a broader range of gating phenotypes than previously anticipated. Our work points towards formerly unknown contributions of NALCN to neuronal excitability and opens avenues for pharmacological targeting.

**Highlights:**

- Function of NALCN requires UNC79, UNC80 and FAM155A
- The complex is permeable to monovalent cations, but is blocked by divalent cations
- The complex displays a constitutively active, voltage-modulated current phenotype
- Positively charged side chains in S4 of NALCN VSD I and II confer voltage sensitivity

## Introduction

Many neurons display a basal Na^+^ conductance at rest that is involved in the regulation of resting membrane potential (RMP), spontaneous firing and pacemaking activity (Raman et al., 2000, Jackson et al., 2004, Khaliq and Bean, 2010). The sodium leak channel (NALCN) contributes to this tonic current in certain types of neurons and plays a critical role in their excitability (Lu et al., 2007, Lu et al., 2010, Flourakis et al., 2015, Shi et al., 2016, Lutas et al., 2016, Philippart and Khaliq, 2018). Consistent with this notion, NALCN knockout is lethal within a day after birth in mice due to disrupted respiratory rhythm (Lu et al., 2007). Furthermore, changes in NALCN expression and/or function have been implicated in other physiological processes such as motor function, pain sensitivity and circadian rhythm in animals (Lear et al., 2005, Yeh et al., 2008, Xie et al., 2013, Flourakis et al., 2015, Gao et al., 2015, Eigenbrod et al., 2019). In humans, accumulating evidence shows that mutations in NALCN cause severe congenital neurodevelopmental disorders (Al-Sayed et al., 2013, Chong et al., 2015, Bramswig et al., 2018), thus underlining its physiological significance.

NALCN represents the sole member of a distinct branch of the four-domain ion channel family, which includes the extensively studied voltage-gated sodium and calcium channels (Na_V_s and Ca_V_s) (Ren, 2011). These channels are made up of four homologous domains (DI–IV), each with six transmembrane segments (S1–S6). Functionally, the voltage-sensing domains (VSDs), formed by S1–S4 of each domain, detect changes in membrane potential and induce the opening or closing of the ion-conducting pore domain, formed by S5 and S6. Unlike Na_V_s and Ca_V_s, however, the fundamental electrophysiological and pharmacological properties of NALCN remain largely unexplored due to poor heterologous expression (Swayne et al., 2009, Senatore et al., 2013, Boone et al., 2014, Egan et al., 2018). In this study, we found that the robust functional expression of NALCN requires UNC79, UNC80 and FAM155A, three ubiquitous neuronal proteins that have been independently shown to interact with NALCN (Yeh et al., 2008, Lu et al., 2009, Lu et al., 2010, Xie et al., 2013). This enabled the first in-depth functional characterisation of the NALCN channel complex and unequivocally defines NALCN as permeable to small monovalent cations, but potently blocked by divalent cations through a direct pore-blocking mechanism. Furthermore, we show that NALCN does not function as a simple Ohmic leak channel, but displays voltage-dependent gating mediated primarily by positive charges in the S4 of DI and II.

## Results

### Co-expression of NALCN, UNC79, UNC80 and FAM155A results in robust currents

Whether NALCN alone can generate leak currents in heterologous expression systems has remained a matter of debate (Lee et al., 1999, Lu et al., 2007, Swayne et al., 2009, Senatore et al., 2013, Boone et al., 2014, Egan et al., 2018). Here, we found that the expression of NALCN alone did not result in detectable responses to a wide range of voltage steps in HEK293 cells or *Xenopus laevis* oocytes (Figure 1A and S1). We thus set out to test if functional expression of NALCN, like that of some of the Ca_V_s (Dolphin, 2016), is dependent on the presence of additional auxiliary proteins. We found that functional expression of NALCN in both heterologous systems requires UNC79, UNC80 and FAM155A (Figure 1A and S1). Furthermore, rat NALCN (Lu et al., 2007), human FAM155B and mouse FAM155A (Xie et al., 2013) orthologs can functionally substitute human NALCN and FAM155A, respectively (Figure S1B). UNC79 and UNC80 are large, potentially disordered proteins (2635 and 3258 aa, respectively) that are highly conserved among animals (Senatore and Spafford, 2013, Cochet-Bissuel et al., 2014). They contain no recognisable functional domains, and their subcellular localisations remain unclear. By contrast, FAM155A (458 aa) contains at least one transmembrane segment and a conserved cysteine-rich domain (CRD) (Pei and Grishin, 2012, Xie et al., 2013). The *C. elegans* and mouse orthologs have previously been suggested to reside in the endoplasmic reticulum (ER) and facilitates NALCN trafficking. For simplicity, we refer the NALCN-UNC79-UNC80-FAM155A combination as the NALCN channel complex henceforth.

**Figure 1.**
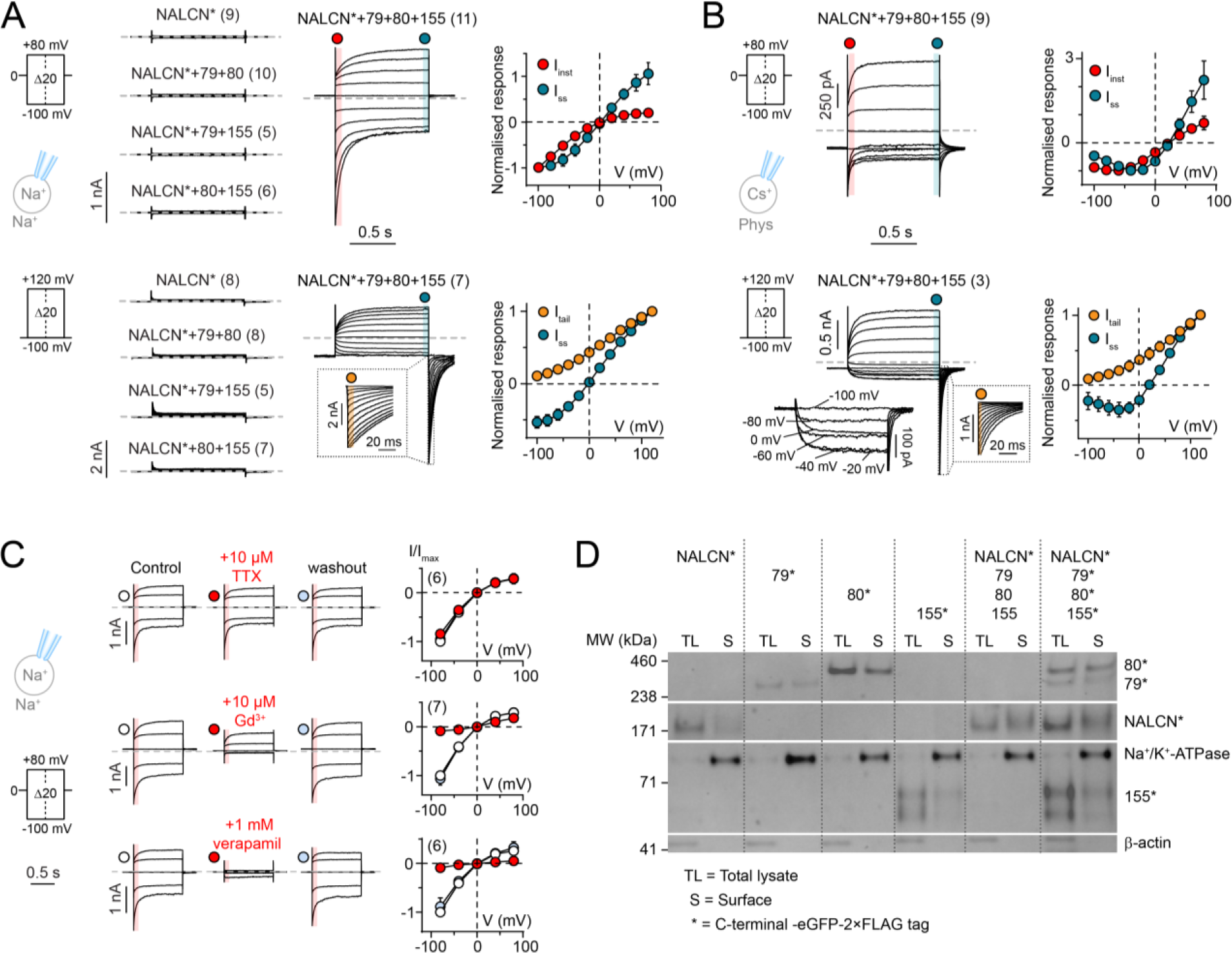
Functional expression of NALCN requires UNC79, UNC80 and FAM155A. (A-B) Whole-cell patch-clamp recordings from HEK cells expressing NALCN-eGFP-2×FLAG (NALCN*) alone, or in different combinations with UNC79 (79), UNC80 (80) and FAM155A (155) under (A) symmetrical Na^+^ and (B) more physiological conditions using voltage-step protocols shown on the left. Normalised I-V plots highlighting the different current components of NALCN*+79+80+155 are shown on the right. The instantaneous current (I_inst_; red) was measured in the beginning of a voltage step change, immediately after the transient current settled. The steady-state current (I_ss_; blue) was measured at the end of a voltage step. Insets show tail currents (I_tail_; orange) immediately after repolarisation to −100 mV. (C) Current responses (left) and I_inst_-V plots (right, normalised to the control current) in the absence and presence of TTX, Gd^3+^ or verapamil under symmetrical Na^+^ condition. Data in A-C are shown as mean ± SD; grey dashed lines indicate 0 nA; numbers in parentheses indicate number of individual cells used for recordings. (D) Western blot of total lysate and surface fraction proteins extracted from HEK cells expressing the indicated constructs (see also Figure S1E).

In patch-clamp experiments, we found that HEK293 cells expressing all components of the NALCN channel complex showed low seal resistances (R_m_∼100 MΩ) after breaking into whole-cell mode (Figure S1D). Despite the low seal resistances, a feature typically associated with non-specific Ohmic leak current in electrophysiological studies (Boone et al., 2014), we observed clear voltage-dependent changes in current in response to both depolarising and hyperpolarising voltage steps, irrespective of the holding potential (HP) (Figures 1A-B, S1A). The gating behaviour of the NALCN complex observed is unique in a few aspects. First, we did not observe full channel closure within the tested voltage range, indicating that the complex is constitutively active (Figure 1A-B). Second, the complex is differentially modulated by voltage, with depolarisation eliciting non-inactivating current and hyperpolarisation eliciting large inward currents that deactivate rapidly but incompletely. Third, the inward component during hyperpolarising steps is much more pronounced under symmetrical Na^+^ condition than under more physiological condition (Figure 1A-B), suggesting that ionic species affect channel function. Fourth, a Na_V_-like, bell-shaped I-V relationship between −100 to 0 mV, with a maximal inward current around −40 mV was detected for the NALCN complex under a more physiological recording condition (Figure 1B). Overall, the gating phenotype is distinct from that of other voltage-gated ion channels (VGICs), which typically populate one or more closed (or inactivated) states upon hyper- or depolarisation.

Next, we investigated if the limited pharmacological profile established for NALCN thus far is preserved when the channel is co-expressed with UNC79, UNC80 and FAM155, i.e. insensitivity to TTX, inhibition by low micromolar concentrations of Gd^3+^ and high micromolar concentrations of the Ca_V_ inhibitor verapamil. We found that TTX application had no effect on current amplitude, while both Gd^3+^ and verapamil showed inhibitory effects (Figure 1C). We also assessed the subcellular localisation of NALCN, UNC79, UNC80 and FAM155A, and found that all four proteins showed membrane localisation, both when expressed alone or together in HEK cells (Figures 1D and S1E). To determine functionally critical regions of these proteins, we expressed full-length NALCN with a series of truncated constructs for UNC79, UNC80 and FAM155A in *Xenopus laevis* oocytes and measured the resulting currents. We found that robust function required the presence of virtually full-length UNC79 and UNC80 proteins, although short truncations were tolerated at the C- and N-terminus, respectively (Figure S2A-B). In the case of FAM155A, the presence of the first putative transmembrane domain and the CRD were absolutely required for function, while deletion of a second putative transmembrane domain was less detrimental (Figure S2C). Together, the data suggest that although NALCN can traffic to the membrane by itself, co-expression with UNC79, UNC80 and FAM155A is a prerequisite for the formation of a functional NALCN channel complex.

### The NALCN channel complex is selective for monovalent cations

To define the ion selectivity profile of the NALCN channel complex, we first determined the current-carrying ions under bionic conditions. We found that current directionality and reversal potentials (E_rev_s) were sensitive to substitution of either extracellular or intracellular Na^+^ with the large cation N-methyl-D-glucamine (NMDG^+^), but not to replacement of extracellular Cl^-^ with the large anion methanesulfonate (MS^-^) (Figure 2A), consistent with permeability for cations and not anions. Next, we assessed the permeability of different cations (Li^+^, K^+^, Cs^+^, TEA^+^, Ca^2+^, Mg^2+^ and Ba^2+^) by having equimolar concentrations of test cations on the extracellular side (150 mM for monovalent cations (X^+^); 110 mM for divalent cations (X^2+^)) and the impermeable NMDG^+^ (150 mM) on the intracellular side. We observed voltage-dependent currents in the presence of extracellular Li^+^, K^+^ or Cs^+^, but not with TEA^+^ or X^2+^ (Ca^2+^, Mg^2+^, Ba^2+^; Figure 2B). The maximal current amplitudes of Li^+^, K^+^ and Cs^+^ elicited at −80 mV, however, were smaller compared to Na^+^ (57±8 %, 89±11 % and 76±11 % of Na^+^ current, respectively). To measure the E_rev_s of the permeable cations, we ran a ramp protocol from −80 to +80 mV. The permeability ratios (P_Na_/P_X_) calculated based on the E_rev_s revealed a permeability sequence of Na^+^≈Li^+^>K^+^>Cs^+^ (Figure 2C). This permeability sequence was altered to Na^+^≈Li^+^≈K^+^>Cs^+^ when the putative selectivity filter (SF) motif of NALCN (EEKE) was mutated to that of Na_V_s (DEKA; Figure 2D).

**Figure 2.**
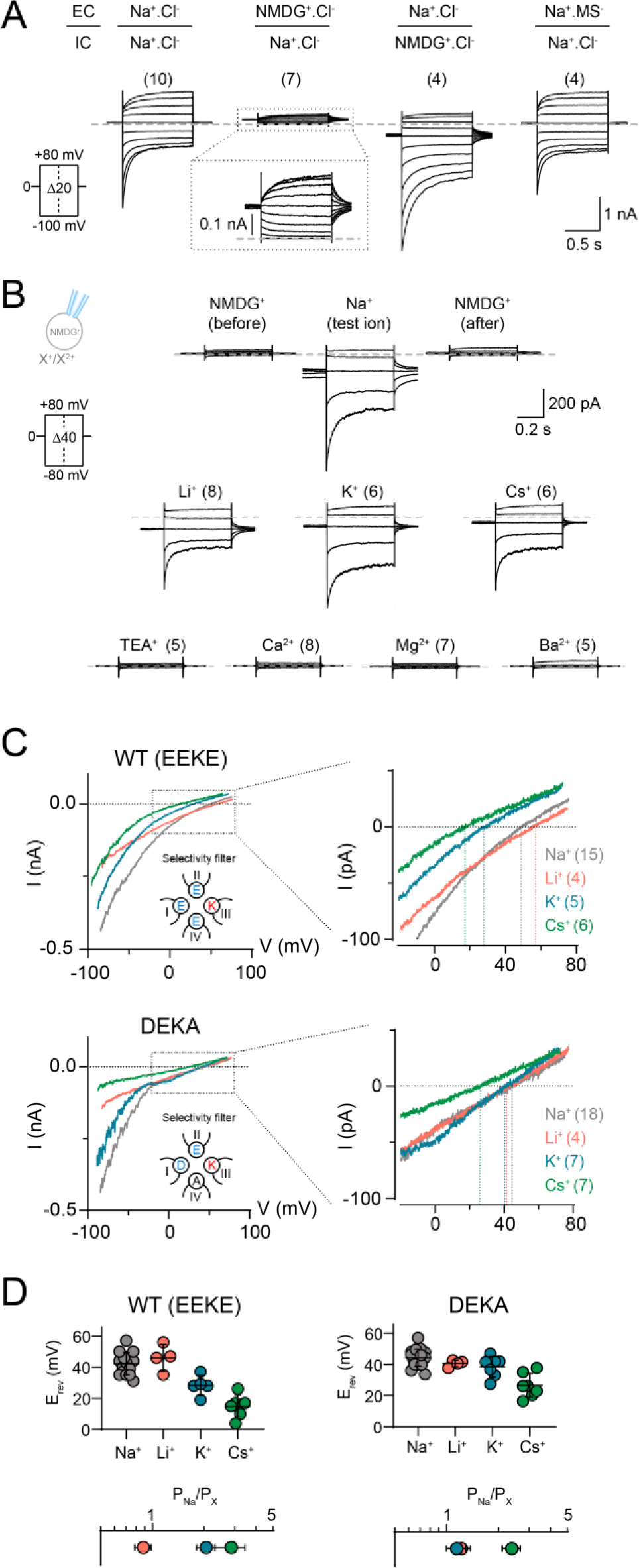
The NALCN channel complex is selective for small monovalent cations. (A) Representative current traces of HEK cells expressing NALCN*+79+80+155 under various bionic conditions in response to the voltage protocol shown. EC=extracellular; IC=intracellular. (B) Representative current traces obtained with intracellular NMDG^+^ and different extracellular mono- or divalent cations using voltage-step protocols shown on the left. (C) The upper panel shows currents obtained with a voltage ramp protocol (−80 to 80 mV) with different test ions for both WT (EEKE SF, top) and the DEKA SF mutant (bottom) using intracellular NMDG^+^ solution; insets show only −20 to +80 mV range. (D) Reversal potentials of different test ions and their permeability relative to Na^+^; data shown as mean ± SD; grey dashed lines indicate 0 nA; numbers in parentheses indicate number of individual cells used for recordings.

### The NALCN pore is blocked by extracellular divalent cations

To investigate if X^2+^ are merely impermeable or directly blocking the channel, we exposed cells to extracellular solutions containing 120 mM Na^+^ in the absence and presence of 1 mM of Ca^2+^, Mg^2+^ or Ba^2+^. We found that all three X^2+^ inhibited Na^+^ currents to varying degrees, with Ca^2+^ being the most potent, followed by Mg^2+^ and Ba^2+^ (83±10 %, 59±11 % and 37±11 % inhibition of I_inst_ respectively; Figure 3A). We also determined the sensitivity of Ca^2+^ inhibition by measuring current responses in the presence of a broad range of extracellular [Ca^2+^] (1 nM to 10 mM) under symmetrical Na^+^ condition. This resulted in an IC_50_ of 319 μM (95 % CI: 192–692 μM) for I_inst_, and 132 μM (95 % CI: 70–350 μM) for I_ss_ (Figure 3B).

**Figure 3.**
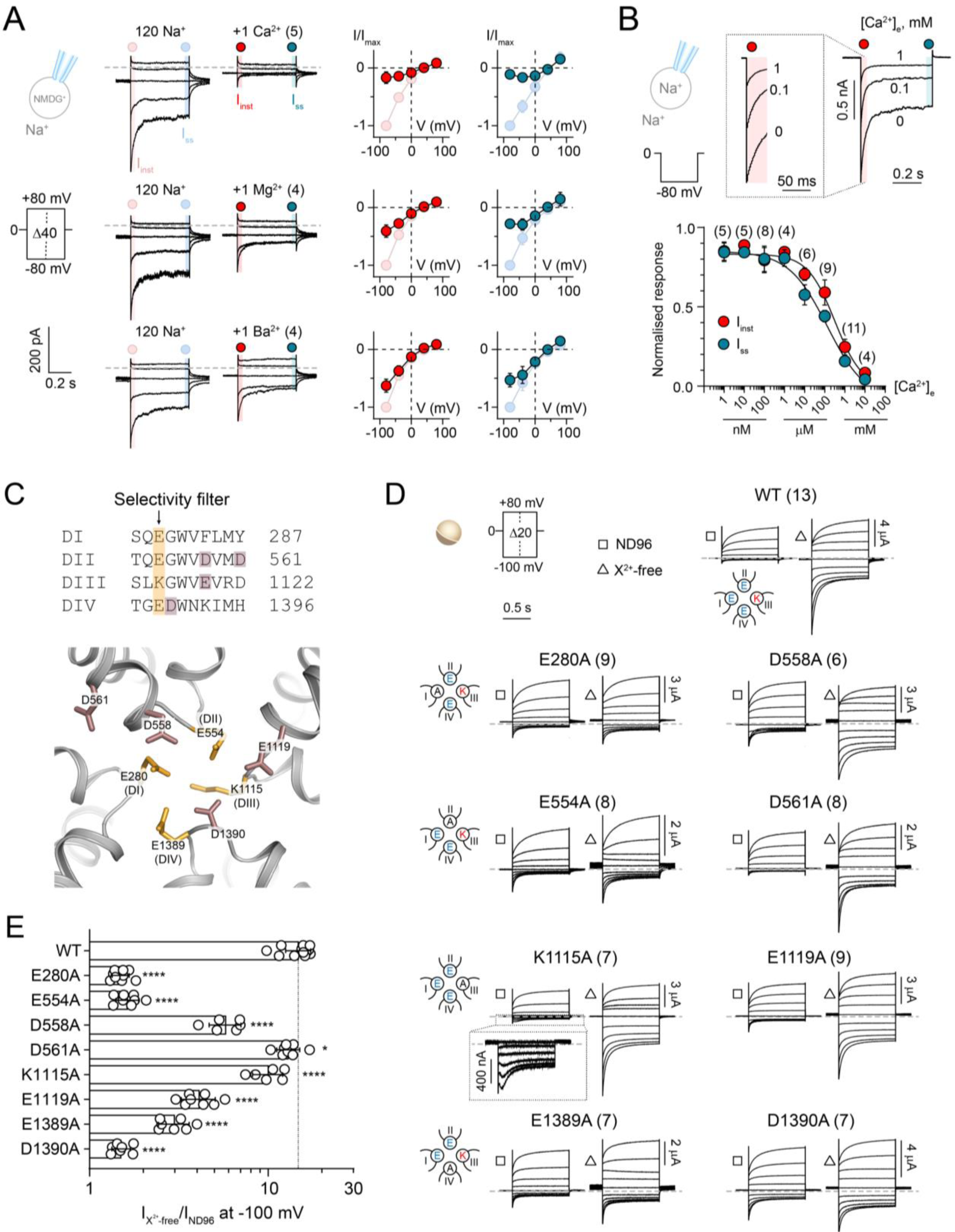
Divalent cation block of the NALCN channel complex is attenuated by mutations of side chains in the putative SF region. (A) Current responses of HEK cells expressing NALCN*+79+80+155 in the absence and presence of 1 mM Ca^2+^, Mg^2+^ or Ba^2+^. Normalised I-V plots illustrate the inhibitory effects of each divalent cation on I_inst_ and I_ss_. (B) Sample traces of a HEK cell expressing NALCN*+79+80+155 exposed to 0, 0.1 and 1 mM Ca^2+^ under symmetrical Na^+^ condition when stepping from 0 to −80 mV. Inset shows I_inst_ on an expanded time scale. IC_50_ graph shows the potency of Ca^2+^ inhibition on both I_inst_ and I_ss_ of NALCN-mediated current. (C) Top, alignment of the selective filter (SF) region from the four homologous domains in NALCN. The putative SF motif EEKE is highlighted in orange and other negatively charged residues selected for charge neutralisation are highlighted in dark pink. Bottom, homology model of the putative SF region of NALCN based on the structure of Ca_V_1.1 (PDB ID: 5GJV). (D) Representative currents from *Xenopus laevis* oocytes expressing WT NALCN or alanine mutants in response to step protocols from +80 to −100 mV (HP=0 mV) in the presence (ND96; 1.8 mM Ca^2+^ and 1 mM Mg^2+^) and absence of divalent cations (X^2+^-free). (E) Fold-increase in inward current elicited at −100 mV for WT NALCN and SF alanine mutants in response to removal of divalent cations. Data are shown as mean ± SD; *, *p*<0.05; ****, *p*<0.0001; One-way ANOVA, Dunnett’s test (against WT); grey dashed lines indicate 0 nA; numbers in parentheses indicate number of individual cells used for recordings. See Figure S3.

We hypothesised that the net negative charge around the putative EEKE SF of NALCN (Figure 3C) influences the sensitivity of X^2+^ block, analogous to what has been demonstrated for Na_V_s and Ca_V_s (Yang et al., 1993, Schlief et al., 1996). To test this notion directly, we substituted eight charged side chains in the putative SF region of NALCN with alanine and compared the effects of removing X^2+^ between wild-type (WT) and mutant channel complexes expressed in *Xenopus laevis* oocytes. While the WT channel complex showed a large increase (15-fold) in inward currents when X^2+^ (1.8 mM Ca^2+^ and 1 mM Mg^2+^ in ND96) were removed (Figure 3D-E), the alanine mutations affected the complex’s sensitivity to X^2+^ removal to varying degrees. E280A, E554A and D1390A showed drastically reduced current increase (1.5-fold), D558A, E1119A and E1389A showed moderately reduced current increase (3-to 6-fold), while D561A and K1115A showed WT-like (>10-fold) sensitivity (Figures 3D-E and S3A). These experiments confirm the direct inhibitory effect of X^2+^ on NALCN and suggest that X^2+^ block the channel by interacting with negatively charged side chains around the putative EEKE SF. The C-terminal tail of NALCN has previously been implicated in an indirect Ca^2+^-sensing mechanism of NALCN that involves the calcium-sensing receptor (Lu et al., 2010). However, we found that C-terminally truncated NALCN variants (Δ1638–1738 and Δ1570–1738) are as sensitive to X^2+^ as the WT channel (Figure S3B), indicating that the direct effects of X^2+^ on NALCN are not dependent on its C terminus.

### Delineating the molecular basis of voltage sensitivity in NALCN

The reduced number of positively charged side chains in the S4 segments of the VSDs of NALCN has previously been suggested to be the cause of its lack of voltage sensitivity (Lu et al., 2007). Here, however, we observed clear voltage-dependence in the NALCN channel complex (Figure 1). To determine the basis of voltage sensitivity, we mutated positively charged side chains in S4 to glutamine. We found that neutralising most, if not all S4 positive charges in VSDI (R146Q, R152Q and R155Q) or VSDII (R481Q, R484Q and K487Q) resulted in channels that did not respond to voltage pulses at HP of 0 mV, but were still sensitive to depolarising pulses from a HP of −100 mV (Figures 4B and S4A-B). In contrast, robust WT-like currents were still elicited in neutralised VSDIII (R989Q, R992Q and R995Q) and IV (R1310Q) mutants (Figure 4B), indicating that the S4 segments in domains III and IV are not essential for the voltage sensitivity of NALCN. Next, we examined the function of single and double charge-neutralising mutations in VSDI and II. All single and double VSDI mutants exhibited WT-like voltage sensitivity and current kinetics (Figure 4C-F), except for R143Q+R146Q and R146Q+R152Q. The R143Q+R146Q mutant exhibits WT-like voltage sensitivity but has significantly larger activation time constants than WT (Figures 4C and E). The R146Q+R152Q mutant, like the triple R146Q+R152Q+R155Q mutant, did not respond to voltage pulses at HP of 0 mV (Figure 4C), but was still sensitive to depolarising pulses from a HP of −100 mV (Figure S4). These results indicate that the presence of two arginine residues in the middle of the VSDI is critical for voltage sensitivity. In VSDII, the single R481Q mutation altered current kinetics, and neutralising an additional VSDII charge on the background of R481Q rendered the channel irresponsive to voltage pulses at HP of 0 mV (Figures 4D-E and S4). Furthermore, R481Q showed significantly slower deactivation time constants compared to WT (Figure 4F). Taken together, the results suggest that R481Q is a critical determinant of voltage sensitivity in NALCN. Supporting evidence for the intrinsic voltage dependence of VSDI originates from the finding that isolated NALCN VSDI generates voltage-dependent currents (Figure S4D), reminiscent of what is observed with the isolated VSD of Shaker potassium channel (Zhao and Blunck, 2016).

**Figure 4.**
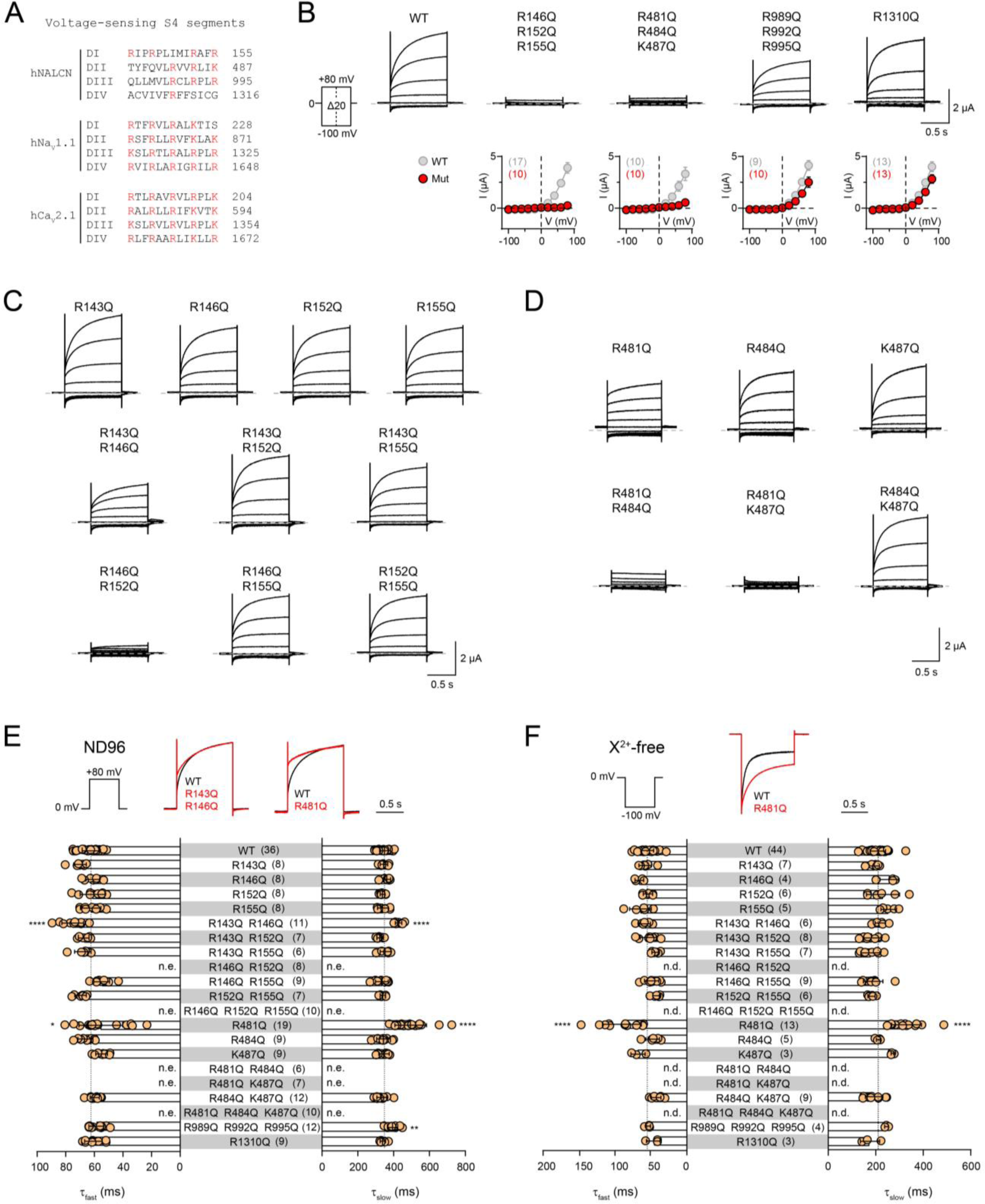
NALCN voltage sensitivity primarily arises from S4 charges in domains I and II. (A) Alignment of the S4 segments of the four homologous domains in hNALCN, hNa_V_1.1 and hCa_V_2.1. Positively charged sides chains are highlighted in red. (B) Current traces and I_ss_-V plots from *Xenopus laevis* oocytes expressing WT and charge-neutralised mutants in S4 of DI-IV using the indicated protocol. (C-D) Representative traces of single and double charge-neutralised mutants in S4 of VSDI (C) and II (D). (E-F) Slow and fast time constants of depolarisation-elicited currents (0 to +80 mV) in ND96 (E) and hyperpolarisation-elicited currents (0 to −100 mV) in X^2+^-free buffer (F) for WT and VSD mutants. Superimposed traces of WT (black) and selected VSD mutants (red) are shown above the bar graphs. Data are shown as mean ± SD; *, *p*<0.05; **, *p*<0.01 ****, *p*<0.0001; One-way ANOVA, Dunnett’s test (against WT); n.e., no effect; n.d.; not determined; grey dashed lines indicate 0 nA; numbers in parentheses indicate number of individual cells used for recordings.

## Discussion

### UNC79, UNC80 and FAM155A are crucial for NALCN function

Two decades have passed since NALCN was first cloned, but functional expression of the channel in heterologous systems is still hampered by issues such as low levels of current (Swayne et al., 2009, Funato et al., 2016, Eigenbrod et al., 2019) and absence of NALCN-specific currents (Lee et al., 1999, Senatore et al., 2013, Boone et al., 2014, Egan et al., 2018). Given the ample evidence supporting overlapping spatiotemporal gene expression profiles for NALCN, UNC79, UNC80 and FAM155A in *Drosophila* and mouse (Ghezzi et al., 2014, Lutas et al., 2016), their functional interdependence in *Drosophila* (Flourakis et al., 2015, Lear et al., 2013), *C. elegans* (Jospin et al., 2007, Humphrey et al., 2007, Yeh et al., 2008, Xie et al., 2013) and mouse (Lu et al., 2009, Lu et al., 2010) and the physical interactions between these proteins *in vitro* and *in vivo* (Lu et al., 2009, Lu et al., 2010, Xie et al., 2013), we set out to test the effect of co-expressing these proteins in different heterologous systems. We found that the robust, reproducible functional expression of NALCN is critically dependent on the presence of UNC79, UNC80 and FAM155A/B (Figure 1A and S1).

UNC79 and UNC80 are widely expressed in the brain, where they form a complex with NALCN and regulate its localisation and function (Yeh et al., 2008, Lu et al., 2010, Lear et al., 2013). While UNC79 is not absolutely required for NALCN activity in mouse hippocampal neurons (NALCN-like currents were observed in the absence of UNC79 (Lu et al., 2010)), both proteins are needed for NALCN function in *Drosophila* (Lear et al., 2013), the neuronal cell line NG108-15 (Bouasse et al., 2019), HEK293 cells and *Xenopus laevis* oocytes (Figures 1 and S1), highlighting potential interspecies differences in the regulatory roles of these proteins. FAM155A is pivotal for the axonal localisation of NALCN in *C. elegans* (Xie et al., 2013). Based on the proposed ER residency of the *C. elegans* and mouse homologues (Xie et al., 2013), it was suggested that FAM155A acts as a chaperone to facilitate NALCN folding and delivery to the cell membrane. By contrast, we were able to detect plasma membrane localisation of the human FAM155A, even when expressed alone (Figure 1D). Whether this discrepancy in subcellular localisation is due to species differences requires further investigation. Overall, our data clearly indicate that NALCN function and not membrane localisation *per se*, is critically dependent on the presence of UNC79, UNC80 and FAM155.

### NALCN is the voltage-sensing, pore-forming subunit of the complex

NALCN was considered to be voltage-insensitive due to the reduced number of voltage-sensing residues in the S4 segments compared to typical VGICs. Here, we found that NALCN, when co-expressed with UNC79, UNC80 and FAM155, exhibits a unique modulation by membrane voltage (Figure 1). This highlights how voltage sensor domains can, depending on the context, contribute to gating in previously unanticipated ways and thus contributes to our understanding of how this widely encountered motif can modulate ion channel function (Bezanilla, 2008). We show that the voltage dependence originates from positively charged residues in the S4 segments of DI and II of NALCN, as charge-neutralising substitutions affected voltage sensitivity (Figure 4). NALCN had been suggested to be a non-selective cation channel that poorly discriminates between X^+^ and X^2+^. This lack of ion selectivity is often attributed to the EEKE SF motif of NALCN, which is a hybrid between the Na^+^-selective DEKA and the Ca^2+^-selective EEEE. Here, we found that NALCN is moderately selective among the permeable X^+^, but does not conduct X^2+^. The X^+^ selectivity profile is abolished when the EEKE motif was mutated to DEKA (Figure 2C). Furthermore, some charge-neutralising mutations around the putative channel pore also affected current rectification and phenotype (Figures 3D and S3A). For instance, the K1115A mutant displays a unique “hooked” phenotype (Figure 3D), consistent with the notion that this side chain lines the ion permeation pathway. Taken together, our results strongly support the idea for NALCN to be the voltage-sensing, pore-forming subunit of the complex.

### The NALCN channel complex reconstitutes the extracellular [Ca^2+^]-sensitive NSCC

A non-selective cation current (NSCC) that is activated by the lowering of extracellular [Ca^2+^] ([Ca^2+^]_e_) has been detected in different types of neurons, such as the chick dorsal root ganglia (Hablitz et al., 1986), mouse hippocampal (Xiong et al., 1997, Lu et al., 2010), neocortical nerve terminals (Smith et al., 2004) and dopaminergic neurons (Philippart and Khaliq, 2018). The NSCC depolarises the RMP, allowing the continuous firing of action potentials. The mechanism(s) underlying this phenomenon are not well understood, but the NALCN channel complex may play a role, as NALCN- and UNC79-knockout neurons are insensitive to drops in [Ca^2+^]_e_ (Lu et al., 2010, Philippart and Khaliq, 2018). This NALCN-mediated effect has been suggested to occur via an indirect mechanism that is initiated and transduced by the G protein-coupled Ca^2+^-sensing receptor (CaSR) (Lu et al., 2010). Here, however, we show that the NALCN channel complex-mediated current, without co-expression of GPCRs, is *directly* blocked by [Ca^2+^]_e_, as alanine mutations of several negatively charged residues in the putative pore of NALCN markedly reduced the sensitivity to [Ca^2+^]_e_ (Figure 3D-E). Furthermore, the NALCN channel complex displays strikingly similar biophysical properties compared to the neuronal NSCC: (1) marked increase in inward Na^+^ current (>10-fold; Figures 1 and 3) following the reduction in [Ca^2+^]_e_, with a similar sensitivity to [Ca^2+^]_e_ (Xiong et al., 1997, Smith et al., 2004, Lu et al., 2010); (2) selective permeability to small X^+^, with a small preference for Na^+^ over K^+^, followed by Cs^+^ (Figure 2) (Xiong et al., 1997); (3) insensitivity to TTX (Raman et al., 2000, Jackson et al., 2004, Khaliq and Bean, 2010) and (4) inhibition by polyvalent cations in the rank order of potency: Gd^3+^>Ca^2+^>Mg^2+^ and Ba^2+^ (Figures 1C and 3A) (Hablitz et al., 1986, Xiong et al., 1997).

### Physiological implications

The contribution of NALCN to the RMP of neurons, and hence their excitability is undisputed. To perform the task as a regulator of the RMP, it is necessary for NALCN to respond dynamically to the potential difference across the plasma membrane. The unique gating behaviour of the NALCN channel complex, along with the modulation by extracellular X^2+^ through a direct pore-blocking mechanism, accentuate its suitability to perform this task. The constitutive activity of the complex implies that a tonic inward Na^+^ conductance will be present even at rest, which may contribute to the depolarisation of the RMP. While this Na^+^ conductance is likely to be small in the presence of normal [Ca^2+^]_e_ and [Mg^2+^]_e_, it is expected to increase considerably following reductions in [Ca^2+^]_e_, such as during repetitive chemical or electrical stimulation, or under pathophysiological conditions (Ren, 2011), thus lowering the threshold for action potential generation. As such, the NALCN complex is predicted to act as an X^2+^ sensor that directly finetunes neuronal excitability in response to fluctuations of [X^2+^]_e_. Furthermore, the differential modulation of NALCN function by voltage means that the NALCN complex undergoes partial deactivation in response to hyperpolarisation, which would allow Na^+^ influx to counter the negative shift in membrane potential. By contrast, during prolonged depolarisation, K^+^ ions would leave the cell, contributing to the repolarisation of the membrane potential. Overall, we postulate that the NALCN complex functions as a thermostat that maintains the RMP at a desired value to determine the intrinsic excitability of a neuron.

## Acknowledgements

We acknowledge the Carlsberg Foundation (CF16-0504; SAP), the Independent Research Fund Denmark (7025-00097A; SAP) and the Lundbeck Foundation (R252-2017-1671; SAP) for financial support. We thank Janne Colding, Cristiana Chirvas and Cristina Bulancea for technical assistance and Drs. Lesley Anson, Harley Kurata, Jianmin Cui and members of the Pless lab for helpful comments on the manuscript.

## Author contributions

H.C.C., M.W. and S.A.P. designed the research. H.C.C., M.W., C.W. and L.P.R. performed the experiments. H.C.C., M.W., C.W. and L.P.R. analysed the data. S.A.P. supervised the project. H.C.C. and S.A.P. wrote the manuscript with input from all authors.

## Supplemental material

### Supplemental Figures

**Figure S1.**
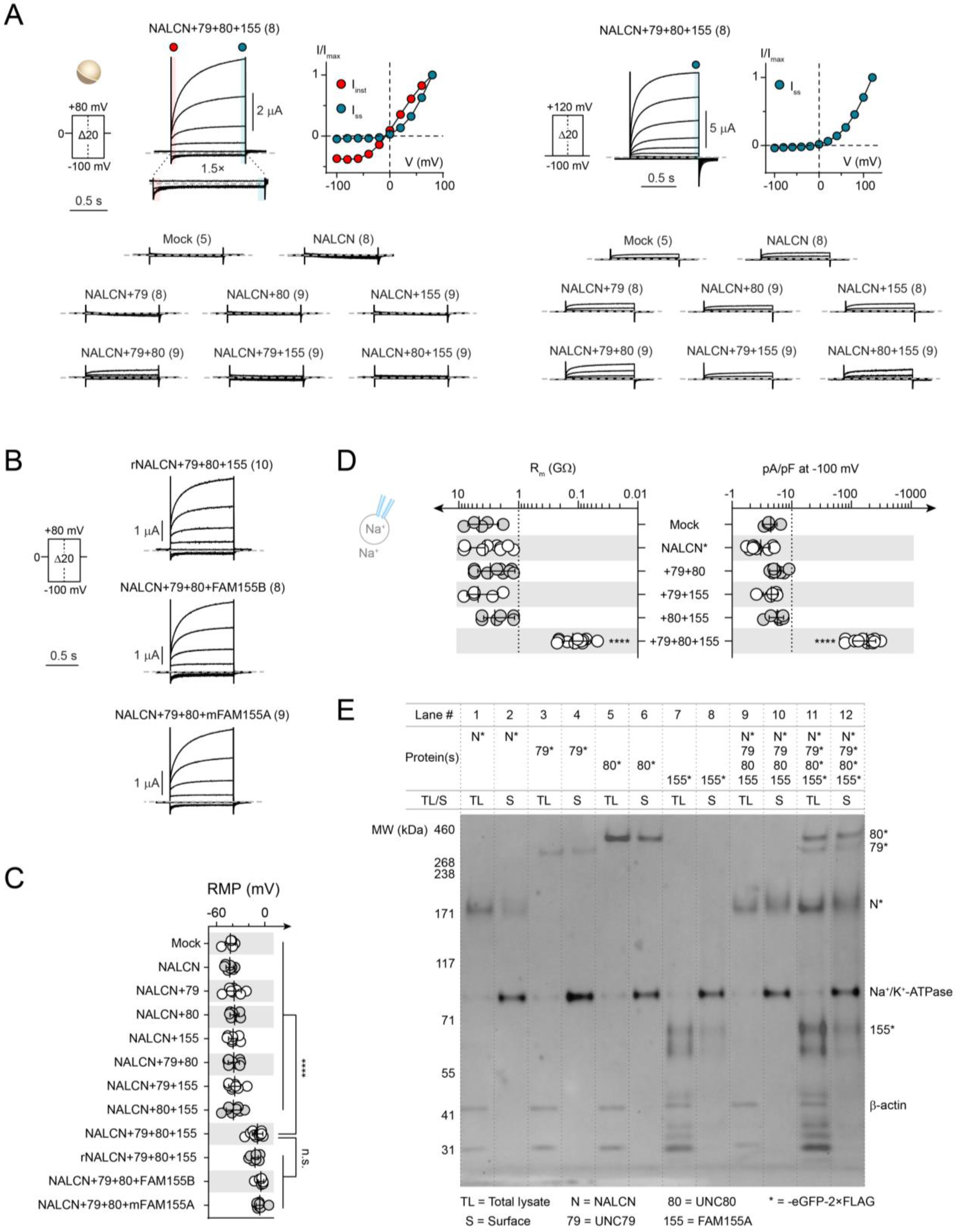
Basic functional and biochemical properties of the NALCN channel complex. (A) Representative current traces from *Xenopus laevis* oocytes expressing NALCN alone, or in different combinations with UNC79 (79), UNC80 (80) and FAM155A (155) using a holding potential of 0 mV and −100 mV (left and right panel, respectively). Normalised I-V plots highlighting the different current components of NALCN+79+80+155 are shown next to the current traces. (B) Representative current traces from *Xenopus laevis* oocytes expressing rat NALCN (rNALCN) with UNC79, UNC80 and FAM155A, NALCN with UNC79, UNC80 and human FAM155B (FAM155B) or mouse FAM155A (mFAM155A) using the indicated protocol. (C) Resting membrane potential (RMP) of *Xenopus laevis* oocytes expressing the indicated combination of constructs. Data are shown as mean ± SD. n.s, not significant; ****, *p*<0.0001; One-way ANOVA, Dunnett’s test (against NALCN+79+80+155). (D) Seal resistances (R_m_) and current densities (pA/pF) for HEK cells expressing indicated combination of constructs under symmetrical Na^+^ condition. Data are shown as mean ± SD. ****, *p*<0.0001; One-way ANOVA, Dunnett’s test (against mock-transfected cells); grey dashed lines indicate 0 nA; numbers in parentheses indicate number of individual cells used for recordings. (E) Uncropped western blot from Figure 1D.

**Figure S2.**
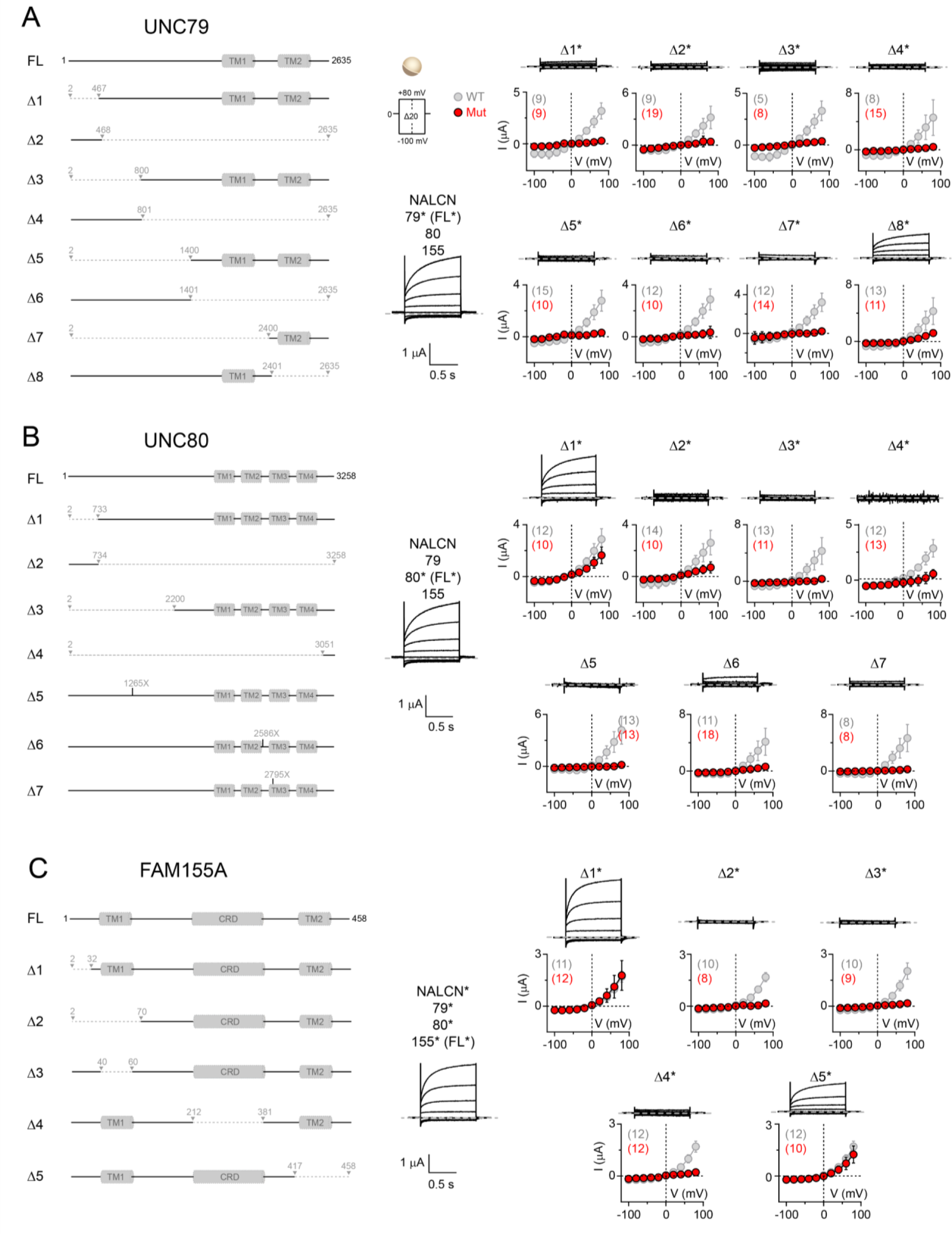
Truncations of UNC79, UNC80 and FAM155A reveal regions critical to NALCN channel complex function. (A-C) Left panels: Schematic depiction of full-length (FL) proteins, along with a series of truncation constructs (ΔX) of UNC79 (A), UNC80 (B) or FAM155A (C), with putative transmembrane (TM) segments shown in grey. The cartoons are made based on the predicted topology of each protein presented on the UniProt database (UNC79: Q9P2D8; UNC80: Q8N2C7; FAM155A: B1AL88). Right panels: Representative currents traces from *Xenopus laevis* oocytes expressing FL or various truncation constructs using the voltage protocol shown in (A). The I_ss_-V plots for WT (grey) and truncation constructs (red) are shown below the individual current traces. Data are shown as mean ± SD; grey dashed lines indicate 0 nA; numbers in parentheses indicate number of individual cells used for recordings.

**Figure S3.**
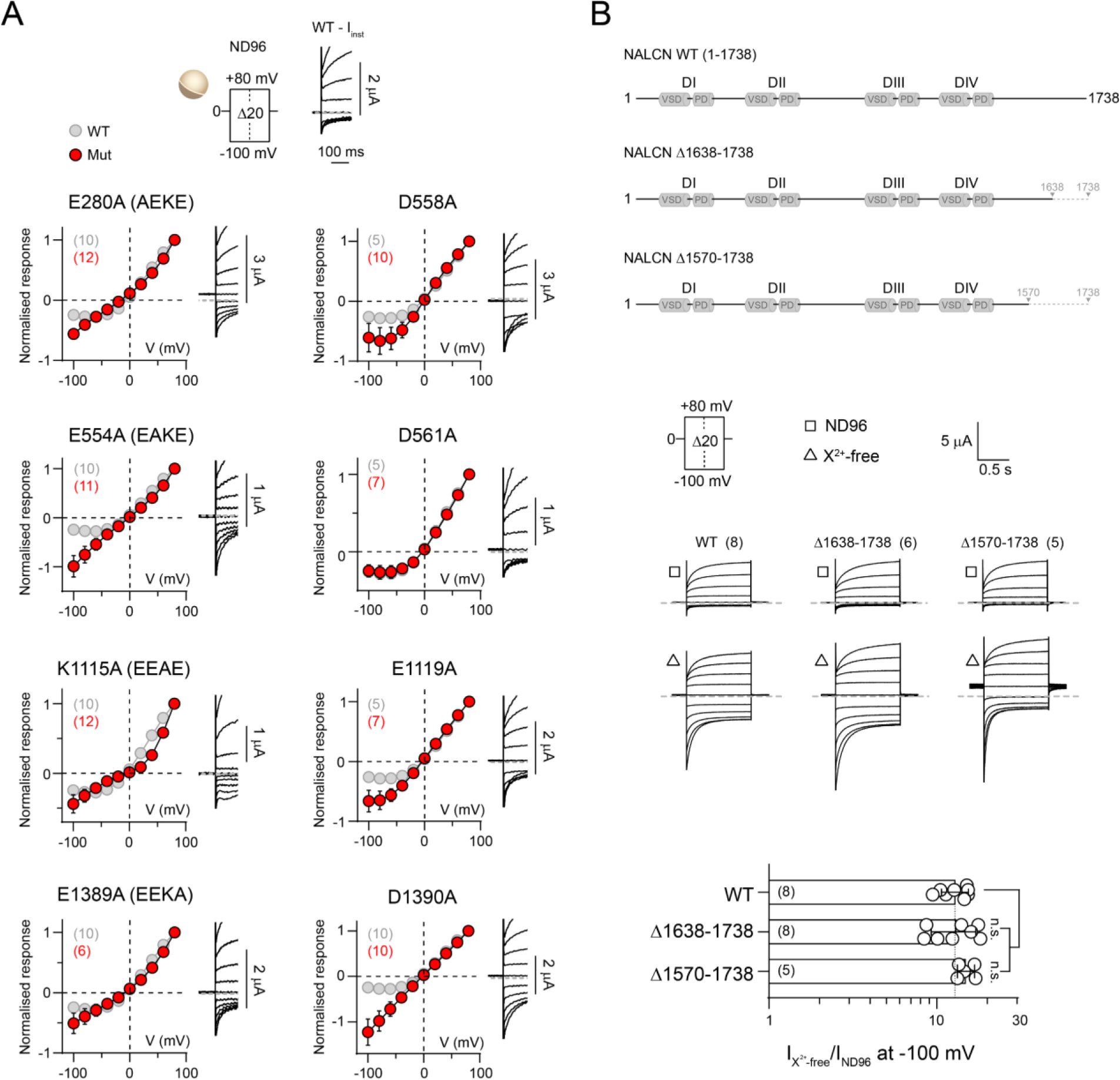
Molecular determinants of X^2+^ sensitivity. (A) I_inst_-V plots (normalised to outward current at +80 mV) from *Xenopus laevis* oocytes expressing WT (grey) and indicated SF alanine mutants (red) recorded in ND96 using the voltage protocol shown on top. Representative current traces highlighting the instantaneous component of WT and SF alanine mutants are shown next to the I-V plots. (B) Top panel: schematic depiction of WT NALCN and C-terminal truncated constructs. Middle panel: representative currents traces from *Xenopus laevis* oocytes expressing WT (1-1738) or C-terminal truncation mutants (Δ1638-1738 and Δ1570-1738) recorded in ND96 (square) and X^2+^-free buffer (triangle) using the indicated voltage protocol. Bottom panel: Fold-increase in inward current elicited at −100 mV for WT NALCN and truncation mutants in response to removal of divalent cations. Data are shown as mean ± SD; n.s., not significant; One-way ANOVA, Dunnett’s test (against WT); grey dashed lines indicate 0 nA; numbers in parentheses indicate number of individual cells used for recordings.

**Figure S4.**
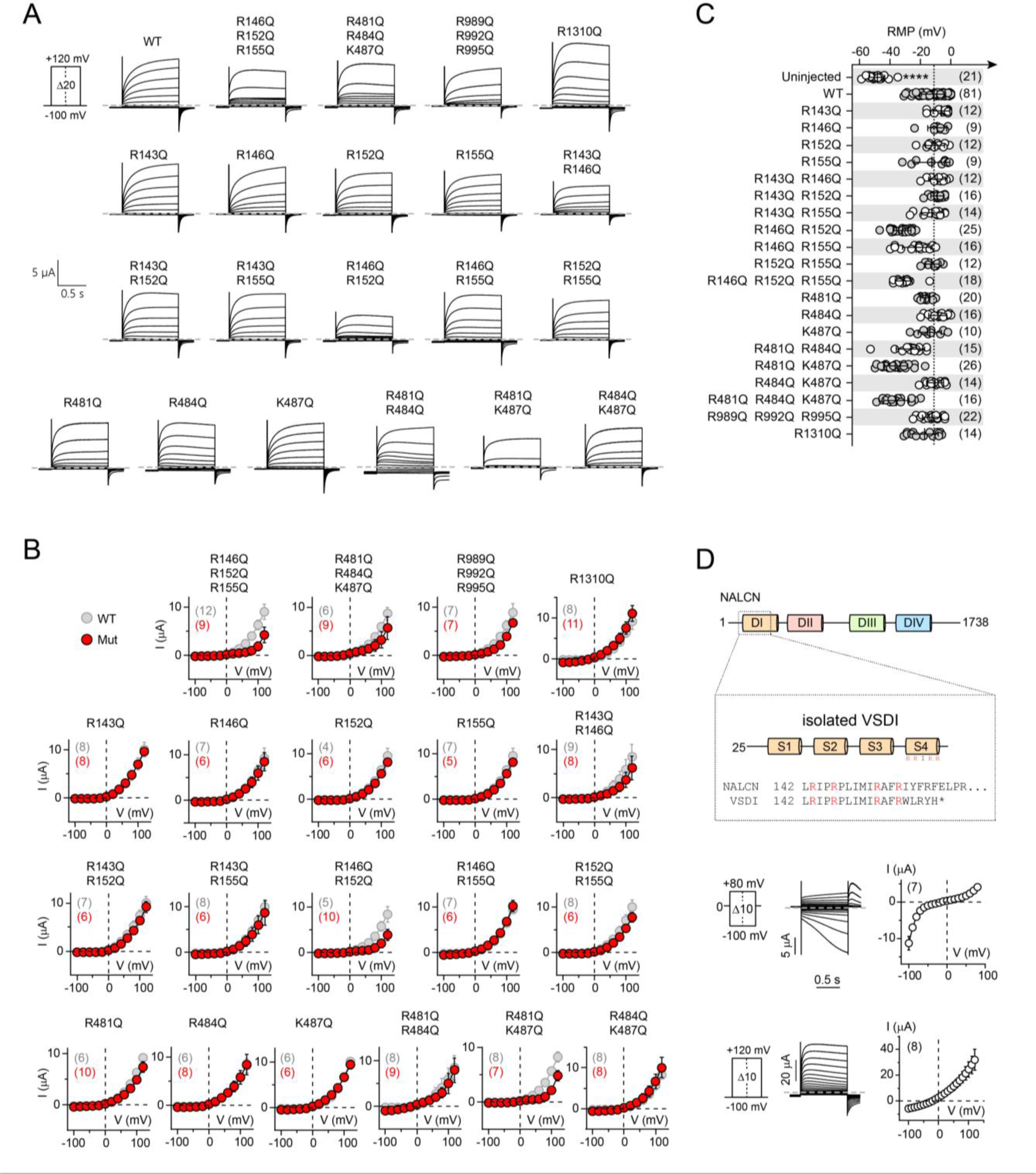
NALCN voltage sensitivity primarily arises from S4 charges in domains I and II. (A-B) Representative traces (A) and I-V plots (B) from *Xenopus laevis* oocytes expressing WT or charge-neutralised VSDI–IV S4 mutants in response to depolarising voltages from −100 to +120 mV. (C) Resting membrane potentials (RMPs) of uninjected, WT- and VSD mutant-expressing *Xenopus laevis* oocytes. (D) Top panel: schematic depiction of full-length NALCN and the isolated VSDI. Bottom panel: representative traces from *Xenopus laevis* oocytes expressing isolated VSDI in response to two different voltage-step protocols. Data are shown as mean ± SD; ****, *p*<0.0001; One-way ANOVA, Dunnett’s test (against uninjected oocytes); grey dashed lines indicate 0 nA; numbers in parentheses indicate number of individual cells used for recordings.

## Materials and Methods

### Molecular biology

Human NALCN*, UNC-79*, UNC-80*, FAM155A* and FAM155B*, and mouse FAM155A* complementary DNAs (cDNAs; * indicates the C-terminal -eGFP-2×FLAG tag) cloned between HindIII and XhoI sites in a modified pCDNA3.1^(+)^ vector containing 3′-*Xenopus* globin UTR and a polyadenylation signal, were generated using custom gene synthesis with codon optimization for *Homo sapiens* (GeneArt, Thermo Fisher Scientific). The tags were removed using the Q5 Site-Directed Mutagenesis Kit (New England Biolabs) to generate wild-type, untagged constructs. The rat NALCN construct was a kind gift from Dr. Dejian Ren at the University of Pennsylvania and was cloned into the same modified pCDNA3.1^(+)^ vector between KpnI and XhoI sites. NALCN pore mutants (Figures 2 and 3) and non-sense mutations bearing UNC80 variants (Figure S2) were generated with site-directed mutagenesis using custom-designed primers (Eurofins Genomics) and PfuUltra II Fusion HS DNA Polymerase (Agilent Technologies). Truncated NALCN, UNC-79*, UNC-80* and FAM155A* constructs (Figure S2 and S3) were generated using the Q5 Site-Directed Mutagenesis Kit. The sequences of purified plasmid DNAs from transformed *E. coli* were verified by Sanger DNA sequencing (Eurofins Genonimcs). For expression in *Xenopus laevis* oocytes, plasmid DNAs were linearized with XbaI restriction enzyme, from which capped RNAs were synthesised using the T7 mMessage mMachine Kit (Ambion). For expression in HEK293T cells, plasmid DNAs purified using the NucleoBond Xtra Midi Plus kit (Macherey-Nagel) were used.

### Two-electrode voltage-clamp electrophysiology

Ovarian lobes were surgically removed from female *Xenopus laevis* frogs anaesthetized in 0.3 % tricaine (procedure approved by the Danish Veterinary and Food Administration; license number: 2014-15-0201-0031), divided into smaller clumps, and defolliculated by shaking at 200 rpm, 37 °C in OR2 (in mM: 82.5 NaCl, 2.5 KCl, 1 MgCl_2_, 5 HEPES; pH 7.4 with NaOH) containing 1 mg/mL of Type I collagenase (Worthington Biochemical Corporation). Healthy-looking stage V-VI oocytes were isolated and injected with 60–70 ng of RNA in a volume of 41–46 nL using a Nanoliter 2010 injector (World Precision Instruments). The NALCN, UNC-79, UNC-80 and FAM155A RNAs were mixed in a ratio of 3:1:1:1. The same ratio was used when NALCN was replaced with rNALCN and when FAM155A was replaced with FAM155B or mFAM155A (Figure S1B). When one or more constructs are excluded (Figure S1A), equivolume of nuclease-free water was added to keep the concentration of each RNA constant across different combinations. Injected cells were incubated in OR3 (50 % (v/v) Leibovitz’s L-15 medium (Gibco), 1 mM glutamine, 250 μg/mL gentamicin, 15 mM HEPES; pH 7.6 with NaOH) at 18 °C, 140 rpm. Four to five days after RNA injection, two-electrode voltage-clamp measurements were performed on oocytes continuously perfused in ND96 recording solution (in mM: 96 NaCl, 2 KCl, 1 MgCl_2_, 1.8 CaCl_2_, 5 HEPES; pH 7.4 with NaOH) or divalent cation-free ND96 (in mM: 96 NaCl, 2 KCl, 5 HEPES; pH 7.4 with NaOH) at room temperature using a Warner OC-725C Oocyte Clamp amplifier (Warner Instrument Corp, USA). Data were acquired using the pCLAMP 10 software (Molecular Devices) and a Digidata 1550 digitizer (Molecular devices), sampled at 10 kHz. Electrical powerline interference was filtered with a Hum Bug 50/60 Hz Noise Eliminator (Quest Scientific). Recording microelectrodes with resistances around 0.2–1.0 MΩ were pulled from borosilicate glass capillaries (Harvard Apparatus) using a P-1000 Flaming/Brown Micropipette Puller System (Sutter Instrument) and were filled with 3 M KCl.

### HEK293T cell culture and transfection

Human embryonic kidney (HEK293T) cells were grown in DMEM (Thermo Fisher Scientific) supplemented with 10 % fetal bovine serum (Thermo Fisher Scientific) and 1% penicillin-streptomycin (10, 000 U/mL, Thermo Fisher Scientific) at 37 °C in a 5 % CO_2_ humidified growth incubator. Cells between passage 6 to 20 were used for experiments and were tested for mycoplasma (Eurofins Genomics) four times throughout this study. For patch clamp experiments, cells reaching 40–60 % confluency in 35 mm cell culture dishes were transiently transfected with constructs of interest using LipoD293 ver. II (tebu-bio) or PEI 25K (Polysciences) 20 to 24 hours before recording. Depending on the transfection reagent used, a total of 2.5 μg (for LipoD293) or 5 μg (for PEI) of cDNAs were used. The NALCN*, UNC-79, UNC-80 and FAM155A cDNAs were mixed in a ratio of 2:1:1:1. When one or more constructs are excluded from the combination, equal amount of empty vector was added to keep the total cDNA amount constant. For mock-transfected cells, empty vector and eGFP were mixed in a 24:1 ratio. For Western blot experiments, cells reaching 20–40 % confluency in 35 mm cell culture dishes were transiently transfected with constructs of interest using LipoD293 two days before the cells were harvested. A total of 0.8 μg of cDNAs was used, with the NALCN*, UNC-79, UNC-80 and FAM155A cDNAs mixed in a ratio of 1:1:1:1.

### Patch Clamp Electrophysiology

On the day of experiment, transfected HEK293T cells were seeded on poly-L-lysine coated glass cover slips at least three hours before recording. Cells were voltage-clamped at room temperature in the whole-cell configuration using an Axopatch 200B amplifier (Molecular Devices). Data were acquired using the pCLAMP 10 software (Molecular Devices) and an Axon Digidata 1550A digitizer (Molecular Devices) at 10 kHz. Patch pipettes were pulled from Kwik-Fil 1.5/1.12 (OD/ID, in mm) borosilicate glass capillaries (World Precision Instruments), and fire polished to resistances around 3.0–8.5 MΩ. A custom-built glass perfusion tool with four adjacent barrels (OD/ID 0.45/1.60, in mm; CM Scientific) controlled by a MXPZT-300R solution switcher (Siskiyou) was used to rapidly exchange extracellular solutions.

For symmetrical Na^+^ condition (Figure 1A), extracellular solution contained (in mM): NaCl (150), HEPES (10), D-(+)-glucose (30), pH 7.4 with NaOH, ∼325 mOsm/L and intracellular solution contained (in mM): NaCl (136), NaF (10), EGTA (5), HEPES (10), Na_2_ATP (2), pH 7.2 with NaOH, ∼309 mOsm/L. For a nire physiological condition (Figure 1B), extracellular solution contained (in mM): NaCl (150), KCl (5), CaCl_2_ (0.5), MgCl_2_ (1.2), HEPES (10), D-(+)-glucose (13), pH 7.4 with NaOH, ∼320 mOsm/L and intracellular solution contained (in mM): CsCl (140), CsF (10), EGTA (5), HEPES (10), Na_2_ATP (2), pH 7.2 with CsOH, ∼304 mOsm/L. For ion selectivity experiments (Figure 2): (1) involving monovalent cations, the extracellular solution contained (in mM): XCl (150), HEPES (10), D-(+)-glucose was added accordingly to achieve osmolarity ∼325 mOsm/L, and the pH was adjusted to 7.4 with XOH, where X indicates the cation of interest; (2) involving divalent cations, the extracellular solution contained (in mM): XCl_2_ (110), HEPES (10), D-(+)-glucose was added accordingly to achieve osmolarity ∼325 mOsm/L, and the pH was adjusted to 7.4 with X(OH)_2_, where X indicates the cation of interest; (3) the intracellular solution contained (in mM): NMDG (150), EGTA (5), HEPES (10), Na_2_ATP (2), D-(+)-glucose (30), pH 7.2 with HCl, ∼310 mOsm/L. For the determination of Ca^2+^ IC_50_ (Figure 3B), extracellular Na^+^ solutions containing 1 nM to 1 mM free Ca^2+^ were made by mixing Ca^2+^-free stock (in mM: NaCl (150), HEPES (10) and EGTA (5), pH 7.4 adjusted with NaOH) and 1 mM free Ca^2+^-stock (in mM: NaCl (150), HEPES (10), EGTA (5), Ca(OH)_2_ (6), pH 7.4 adjust with NaOH) at the ratio calculated according to the WEBMAXC STANDARD calculator (https://web.stanford.edu/~cpatton/webmaxcS.htm). Solution containing 10 mM Ca^2+^ was prepared by adding CaCl_2_ from a 1 M stock solution. D-(+)-glucose was added accordingly to all extracellular solutions to achieve osmolarity ∼325 mOsm/L. The intracellular solution contained (in mM): NaCl (136), NaF (10), EGTA (5), HEPES (10), Na_2_ATP (2), pH 7.2 with NaOH, ∼309 mOsm/L.

To prevent non-specific leaks from affecting the accuracy of our results, we routinely checked for loose seals by exposing cells to NMDG-only extracellular solution before and/or after experiments (Figure 2B). Cells that showed steady-state inward current >10 pA at −80 mV in the absence of permeable ions were discarded. Voltage ramp protocol used to determine E_rev_s (Figure 2C) is as follows: whole-cell mode was first achieved in symmetrical NMDG^+^ solutions, and cells were exposed to NaCl solution for 200 ms before a 1-s voltage step (0 to −80 mV) was applied to obtain a steady-state current; at the end of the voltage step, a 200-ms ramp (−80 to +80 mV) was run before returning to 0 mV; cells were allowed to rest in NDMG solution for ∼10 s before the same protocol was run in a solution containing a different ion of interest.

### Cell surface biotinylation and Western Blots

Cell surface proteins were purified using the Pierce Cell Surface Protein Isolation Kit (Thermo Fisher Scientific) based on manufacturer instructions with a few modifications: (1) washing was performed with ice-cold PBS-CM (in mM: 137 NaCl, 2.7 KCl, 10 Na_2_HPO_4_, 1.8 KH_2_PO_4_, 0.1 CaCl_2_, 1 MgCl_2_), (2) biotin was dissolved in ice-cold PBS-CM to a final concentration of 1.25 mg/mL; and (3) the quenching buffer consisted of PBS-CM, supplemented with 200 mM glycine. Cells were lysed in lysis buffer (100 mM Tris-HCl, 150 mM NaCl, pH 7.4, 0.1 % SDS, 1 % Triton-X-100, 1:100 Halt Protease Inhibitor Cocktail (Thermo Fisher Scientific)) with gentle agitation on ice for 30 min. Denatured samples from both the total lysates and surface fractions were separated on NuPAGE 3-8 % Tris-Acetate protein gels (Thermo Fisher Scientific) at 200 V for 40 min. The HiMark Pre-Stained Protein Standard (30–460 kDa; Thermo Fisher Scientific) was used as protein molecular weight reference. Samples were transferred onto an Invitrolon PVDF membrane activated with 10 % methanol using the iBlot 2 Dry Blotting System (Thermo Fisher Scientific). The membranes were then incubated in blocking buffer (1×TBS supplemented with 4.5 g/L Fish gelatin, 1 g/L casein, 0.02 % sodium azide) for 1 hr at RT before they were incubated with the primary antibodies (rabbit anti-FLAG (701629; Invitrogen), mouse anti-β-actin (sc-47778; Santa Cruz Biotechnology), mouse anti-Na^+^/K^+^-ATPase (05-369; EMD Merck Millipore)) at 4 °C overnight. Next, the membranes were washed 5×2 min with TBST (20 mM Tris-HCl, 150 mM NaCl, 0.1 % Tween 20, pH 7.5) and incubated with the secondary antibodies (IRDye 800CW goat-anti-rabbit (925-32211; LI-COR Biosciences) and IRDye 680RD goat-anti-mouse (926-68070; LI-COR Biosciences)) for 1 hr at RT in the dark. Finally, the membranes were washed 5×2 min in TBST before being imaged using a PXi gel imaging station (Syngene).

### Data analysis

Raw current traces were generally filtered at 500–800 Hz (8-pole Bessel low-pass filter) before data analysis. Current traces were subjected to data reduction (substitute average by a factor of 5) for illustration. Data were presented as mean ± standard deviation (SD). Statistical comparisons were performed using GraphPad Prism (version 8.1, GraphPad Software) and the specific tests used were mentioned in text where relevant. For ion selectivity experiments (Figure 2C), liquid junction potential (LJP) was measured with reference to the NMDG intracellular solution and corrected post recording (Neher, 1992). The measured LJP values for extracellular NaCl, LiCl, KCl and CsCl solutions were 4.5, 2.6, 8.4 and 7.4 mV respectively. Relative ion permeabilities (P_Na_/P_X_) were calculated using the Goldman-Hodgkin-Katz equation P_Na_/P_X_=exp(F(E_rev(Na)_–E_rev(X)_)/RT), where F=Faraday’s constant, R=gas constant, T=274 K, and E_rev_s were measured from the −80 to +80 mV ramp (corrected for LJP). The homology model of human NALCN was generated using the Phyre2 Protein Fold Recognition Server; the model based on the Cav1.1 channel (PDB 5GJV) was selected for the presentation shown in Figure 3C.

## Protein Sequences

### >hNALCN

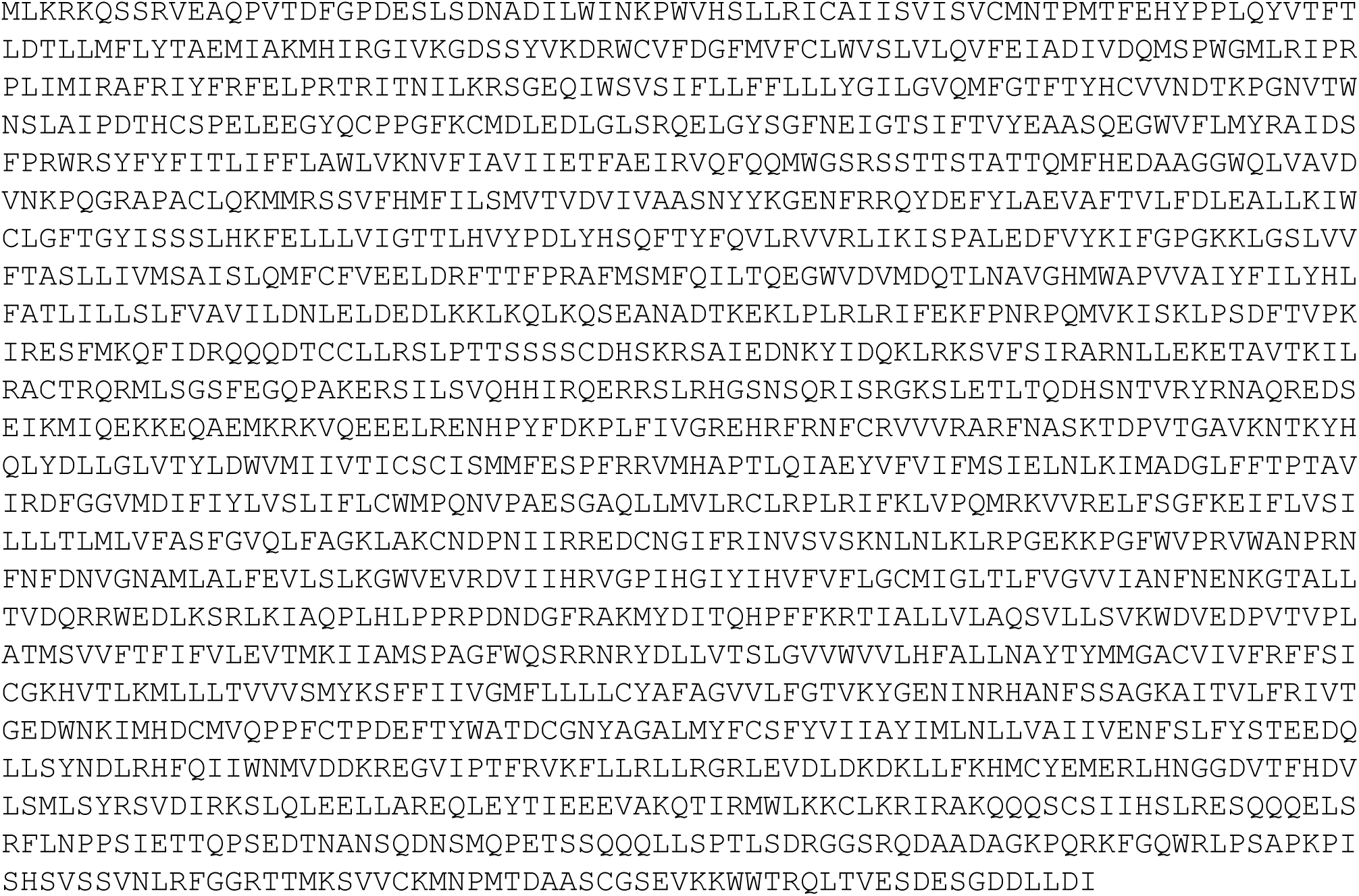

### >rNALCN

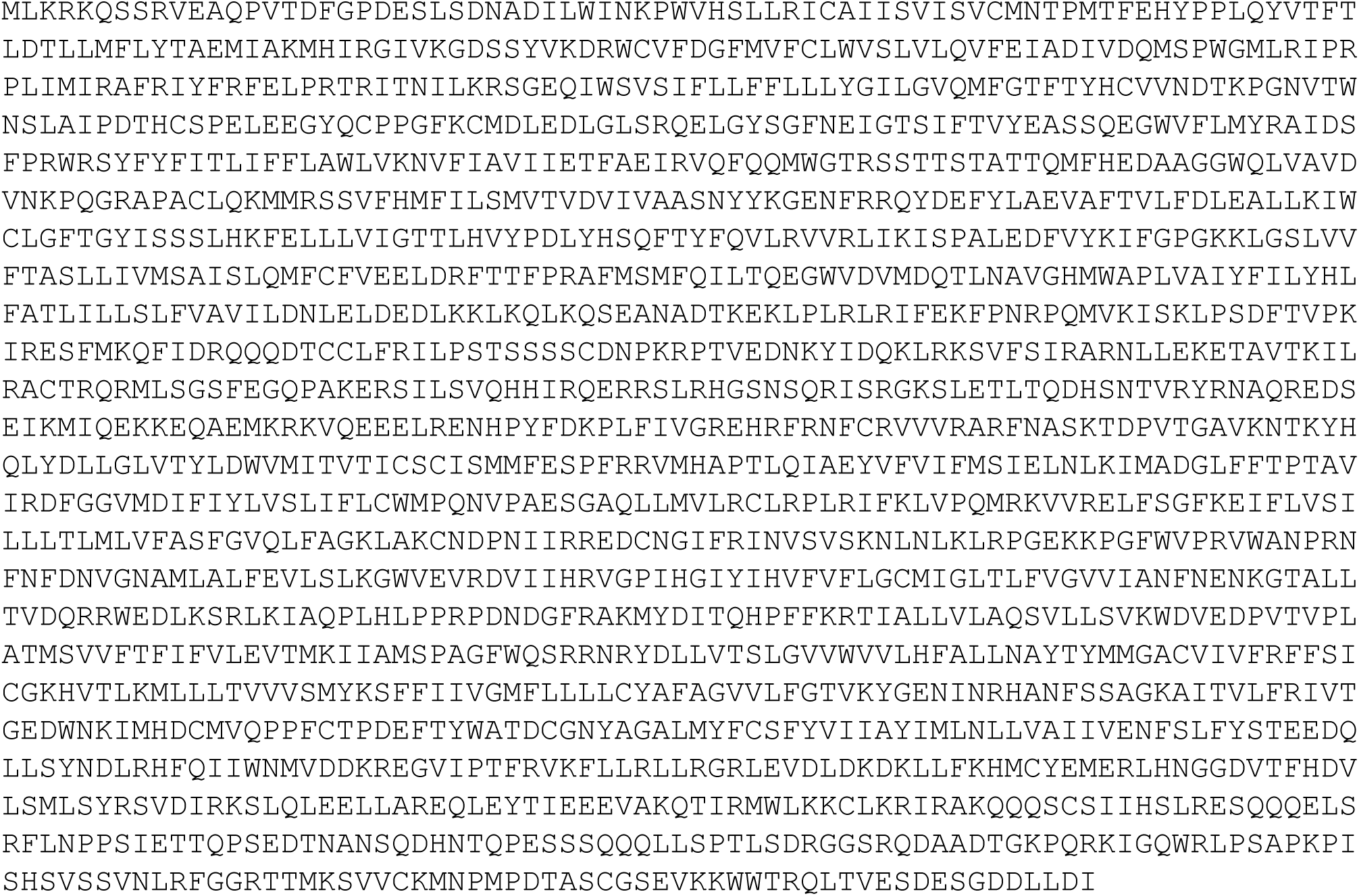

### >hUNC79

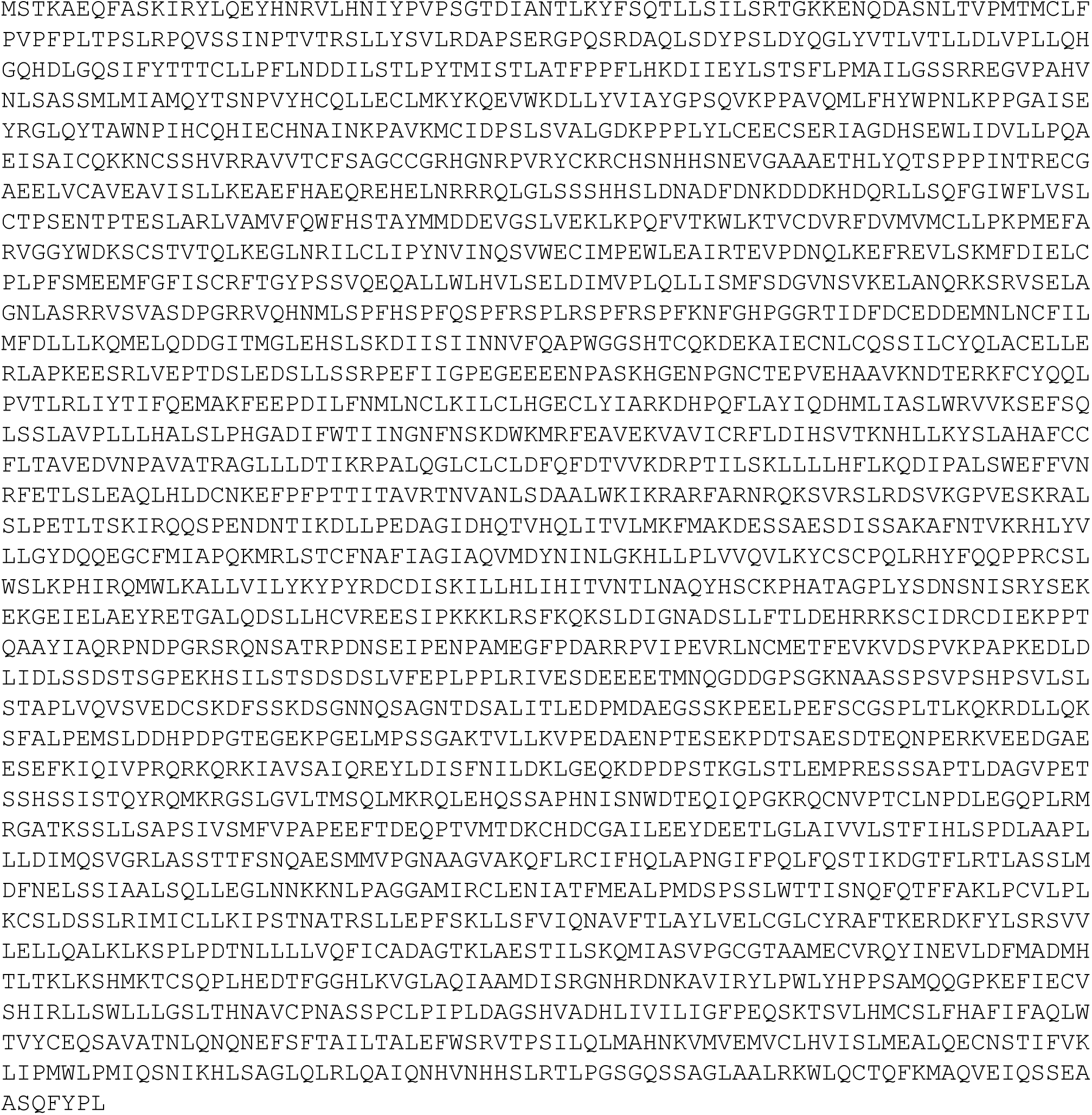

### >hUNC80

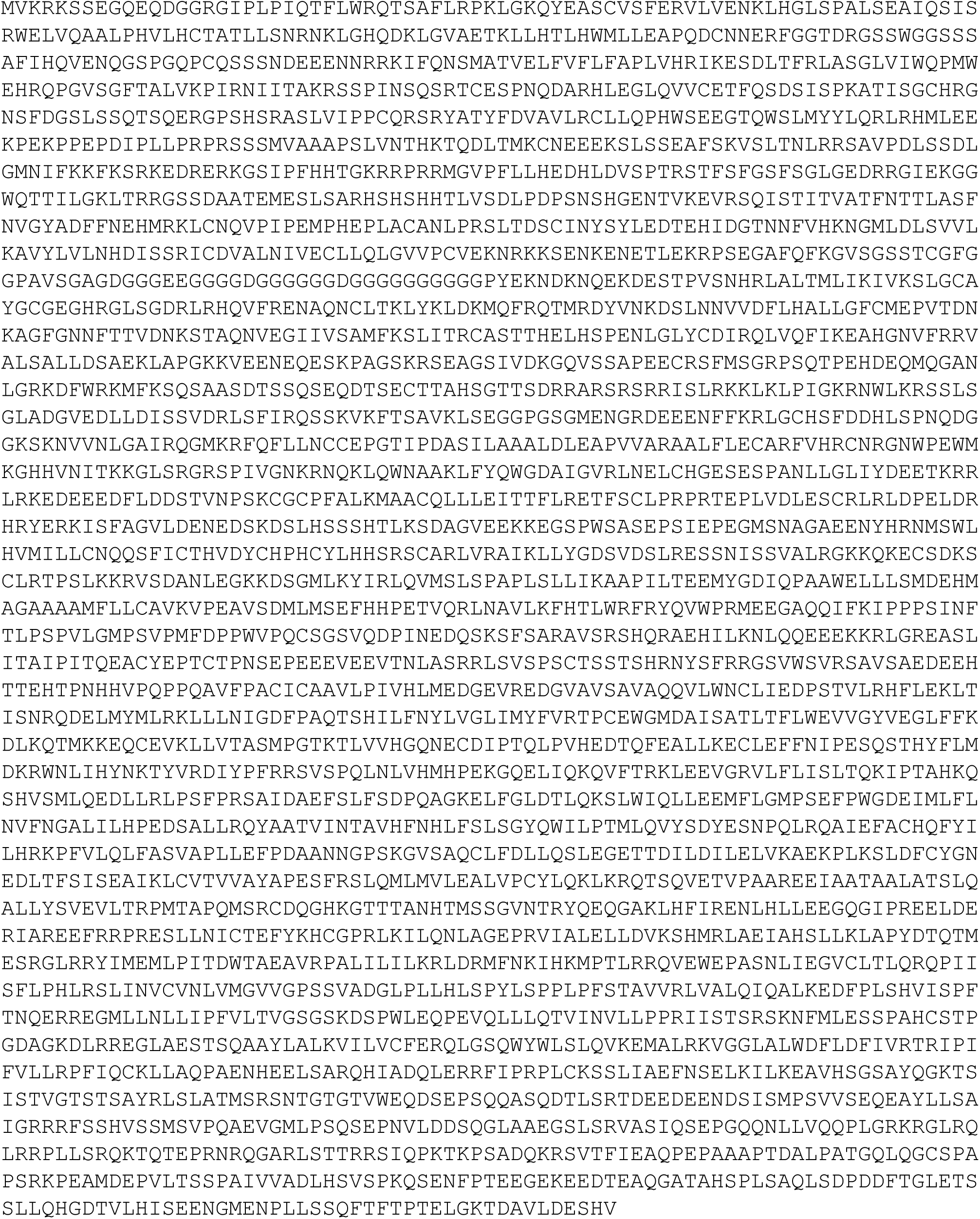

### >hFAM155A

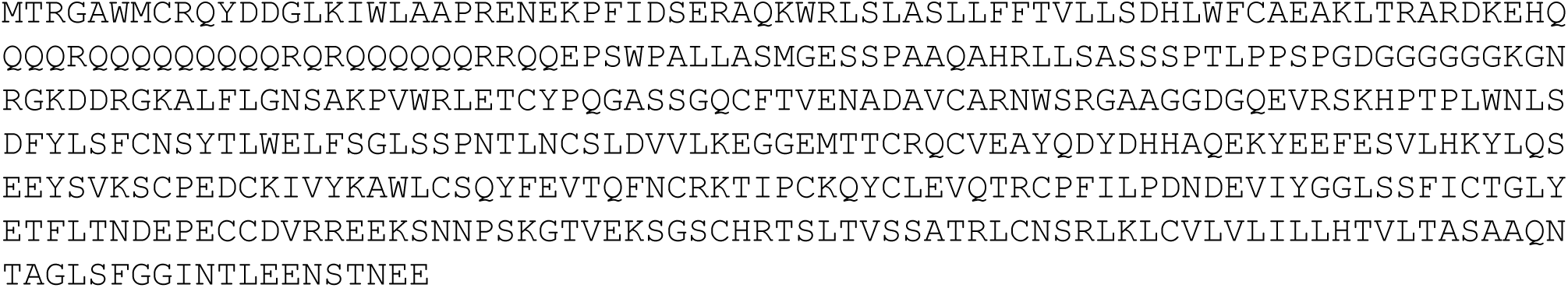

### >hFAM155B

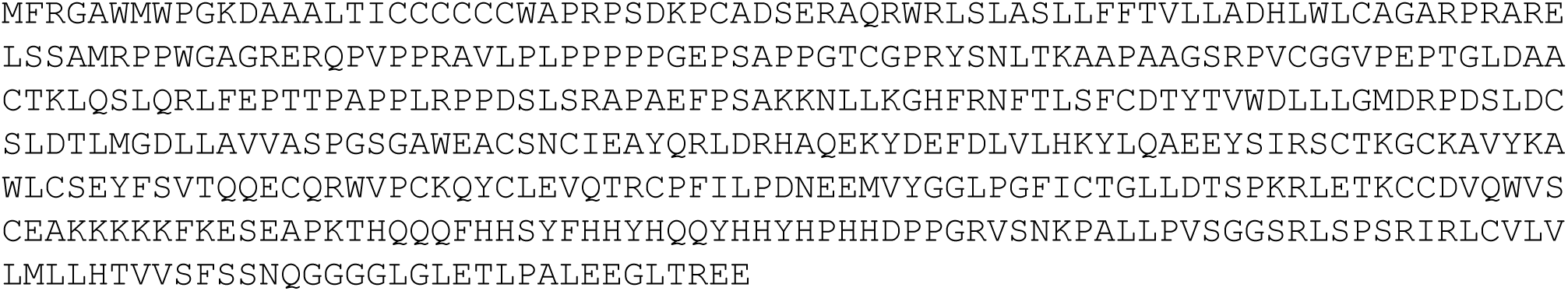

### >mFAM155A

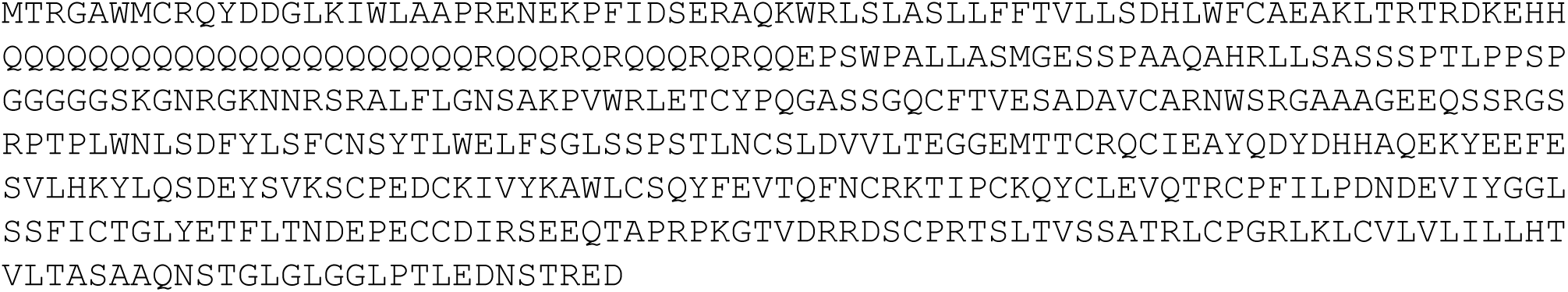

### >isolated VSD1 of NALCN

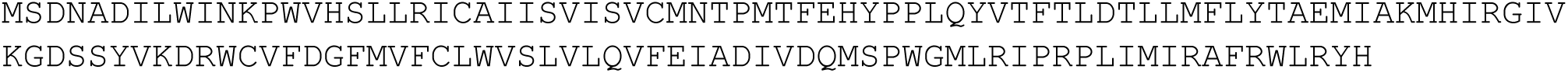

### >C-term_-eGFP-2×FLAG_tag

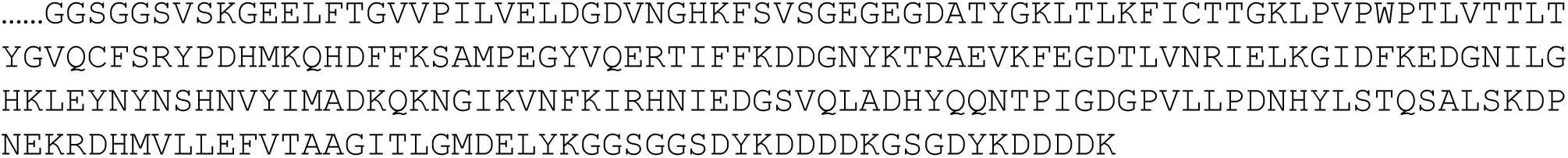

